# Vaginal lactobacilli produce anti-inflammatory β-carboline compounds

**DOI:** 10.1101/2024.02.19.580919

**Authors:** Cecilia Webber, Virginia Glick, Morgan Martin, Cecilia Kim, Maryam Ahmad, Lauren Simmons, Sunghee Bang, Michael Chao, Nicole Howard, Sarah Fortune, Lalit Beura, Seo Yoon Lee, Jon Clardy, Ki Hyun Kim, Smita Gopinath

## Abstract

The vaginal microbiome is a lactobacilli-dominant, human-adapted community consistently found in people around the world. Presence of lactobacilli-dominant vaginal microbial communities, apart from *Lactobacillus iners,* is associated with reduced vaginal inflammation and reduced levels of pro-inflammatory cytokines. Loss of lactobacilli-dominance is associated with inflammatory conditions such as bacterial vaginosis. It remains unclear if health-associated lactobacilli actively promote the anti-inflammatory immune homeostasis of the vaginal mucosa and if so, by which mechanisms. We have identified that *Lactobacillus crispatus,* a key vaginal bacterial species, secretes a family of β-carboline compounds with anti-inflammatory activity. These compounds suppress NFκB and interferon signaling downstream of multiple pattern recognition receptors in human macrophages and significantly dampen type I interferon receptor activation. Topical vaginal application of an anti-inflammatory β-carboline compound, perlolyrine, was sufficient to significantly reduce vaginal inflammation in a mouse model of genital herpes infection. Together, we identify a key family of compounds by which vaginal lactobacilli mediate host immune homeostasis and could highlight a new therapeutic avenue for vaginal inflammation.

## Introduction

The microbiome modulates the immune homeostasis of the mucosa in a variety of ways including production of small molecule effectors. These mechanisms have been under studied in mucosal surfaces beyond the gut. The bacterial communities of the vaginal microbiome can be broadly divided into two iterations – lactobacilli-dominant, low-diversity communities or lactobacilli-low, high-diversity communities (France et al., 2022). Clinical profiling studies have established a robust association between lactobacilli-dominant communities and health including reduced acquisition of sexually transmitted infections (Armstrong and Kaul, 2021; Gosmann et al., 2017), reduced risk of preterm birth (Fettweis et al., 2019) and reduced vaginal inflammation, including production of pro-inflammatory cytokines (Abbe and Mitchell, 2023; Anton et al., 2018). Loss of vaginal lactobacilli is associated with bacterial vaginosis, which in turn increases risk of sexually transmitted infections (Atashili et al., 2008; Bautista et al., 2017) including herpes virus infection (Esber et al., 2015). Research to date has focused on the pathogenesis of non-lactobacilli, BV-associated bacteria (Anton et al., 2022; Joseph et al., 2023). We hypothesized that vaginal lactobacilli actively modulate the immune environment, potentially via secretion of immunomodulatory effectors.

The vaginal mucosal epithelium is primarily composed of epithelial cells however recruited and resident immune cells can have a large impact on tissue level immunity. We previously found that activating vaginal mucosal dendritic cells which formed less than 1% of the total cell population was sufficient to protect against vaginal viral infection (Gopinath et al., 2018). While the effect of vaginal bacteria on epithelial cell cytokine responses has been studied (Doerflinger et al., 2014; Anton et al., 2022, 2018), we do not understand how vaginal bacteria affect immune cells. We hypothesize that vaginal lactobacilli modulate the vaginal immune environment via secretion of effector molecules. The pro-inflammatory cytokines associated with lactobacilli-low communities, vaginitis and bacterial vaginosis include IL-1β, IL-6 and TNFα, many of which are driven by NFκB activation (Anahtar et al., 2018; Abbe and Mitchell, 2023). Likewise, type I interferon (IFN) can also drive pathogenic inflammatory responses in the vaginal mucosa (Lebratti et al., 2021). The effect of vaginal lactobacilli on NFκB and IFN signaling in immune cells is unknown.

There are four main vaginal lactobacilli species found in people - *L. crispatus, L. jensenii*. *L. gasseri* and *L. iners*. Of these species, *L. iners* dominance is associated with transition to loss of lactobacilli and establishment of BV-associated high-diversity communities (Ravel et al., 2013; Gajer et al., 2012). A recent systematic review found that *L. iners* dominance alone is linked to increased vaginal inflammation, bacterial vaginosis, and increased susceptibility to sexually transmitted infections (Carter et al., 2023). *L. iners* has the smallest genome of the vaginal lactobacilli and is known to only produce the L-lactic acid enantiomer in contrast to other lactobacilli which produce both L- and D-lactic acid (France et al., 2016). In this study we use both *L. iners* and the gut lactobacillus *L. reuteri* to study the immunomodulatory effects of *L. crispatus*, the most commonly occurring and health-associated vaginal lactobacillus species. We describe a family of small molecule effectors produced by vaginal lactobacilli, that suppress pro-inflammatory NFκB and IFN signaling in human macrophages, marking the first discovery of vaginal lactobacilli effectors since lactic acid.

## Results

To identify vaginal lactobacilli-secreted immunomodulators, we screened a panel of anti-inflammatory *L. crispatus* and transition-associated *L. iners* strains as well as an *L. reuteri* strain for secreted anti-inflammatory effectors. Bacterial strains were grown from single colony, cultured for 24 hours, further sub-cultured for 48 hours and cell-free supernatant collected. We used a dual reporter THP1 human macrophage-monocyte cell line that allowed us to detect both NFκB and type I interferon (IFN) activity (Fig. 1A). We detected significant suppression of basal NFκB activity compared to media controls in cells treated with cell-free supernatant from *L. crispatus* but not *L. iners* strains (Fig. 1B). Likewise, we saw suppression of interferon signaling in the majority of *L. crispatus* but not *L. iners* - treated cells (Fig. 1C). We confirmed that both *L. crispatus* and *L. iners* reached maximum growth as measured by absorbance within the 48 hour time period (Fig. S1A, B) suggesting that differences in anti-inflammatory activity were unlikely to be attributable to robust growth of *L. crispatus* over *L. iners* strains. Since basal IFN levels were low, we next asked if *L. crispatus* bacterial supernatant could suppress interferon signaling in macrophages that had already been activated. To test this, we stimulated THP1s with LPS and then treated with bacterial supernatant from an *L. crispatus* strain that was a relatively weak suppresser (strain MV-1A-US, HM-636, Fig. 1C) and a candidate *L. iners* strain (strain LEAF-2052A-d, HM-706). *L. crispatus* supernatant was sufficient to suppress LPS-activated IFN signaling to levels below those observed in untreated control cells (Fig. 1D). To expand our investigation to other vaginal lactobacilli strains and other pattern recognition receptors, we tested 12 *L. gasseri* strains, 5 *L. jensenii* strains and 3 additional *L. crispatus* strains and found all suppressed NFκB signaling in macrophages treated with Pam3CSK4, a TLR2 ligand (Fig. S2A). To extend our analyses to endosomal TLRs, we also added supernatant from *L. crispatus* and *L. iners* strains to THP1 cells treated with aminoglycosides, a compound we had previously identified as activating TLR3-mediated IFN signaling (Gopinath et al., 2018, 2020). As with TLR4, we found that *L. crispatus* but not *L. iners* strains suppressed aminoglycoside-induced IFN signaling (Fig. S2B). *L. iners* is unique amongst lactobacilli in being unable to grow in MRS media and our initial screen (Fig. 1B-D) used *L. iners* grown in NYC III media. However, a recent study showed that *L. iners* could be grown in cysteine-supplemented MRS media (Bloom et al., 2022), so we confirmed that cell-free supernatant from *L. iners* grown in cysteine-supplemented MRS media still failed to suppress TLR4 activation (Fig. S2C). We have identified that multiple vaginal lactobacilli species, except for *L. iners,* produce anti-inflammatory immunomodulatory compounds that can suppress inflammatory signaling downstream of cell-surface (TLR2,4) and endosomal (TLR3) pattern recognition receptors.

**Figure 1:**
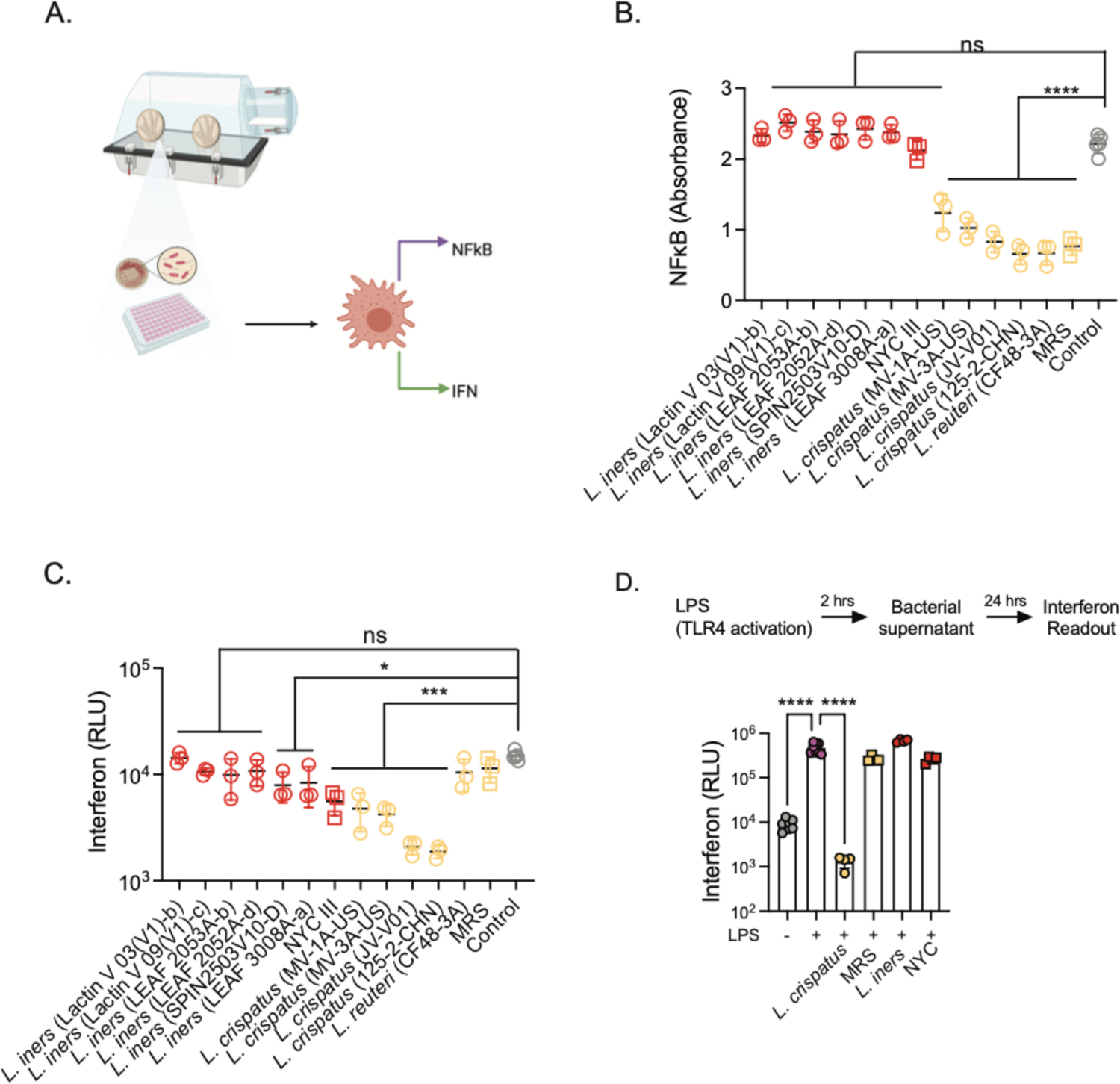
Vaginal lactobacilli-secreted effectors suppress inflammatory signaling in human monocyte macrophage cell line. Indicated bacterial strains were grown from single colonies for 28 hours in NYC III (*L. iners* strains only) or MRS media (all other strains) and filtered-cell-free supernatant added to THP1 reporter monocytes at 5% v/v (A). NFκB (B) and interferon (C) responses were read 8 hours after addition. In (D) THP1 cells were activated with 500 ng/ml LPS and then treated with a candidate *L. crispatus* strain (MV-1A-US) and a candidate *L. iners* strain (LEAF 2052A-d). Samples were compared using a one-way ANOVA with Dunnett’s correction for multiple comparisons. Data in (B) and (C) are representative of 4 experiments with 3 biological replicates per treatment condition. Data in (D) are representative of 2 experiments with 4 biological replicates per treatment condition.

We chose a candidate *L. crispatus* strain (MV-3A-US, HM-636) to perform follow-up analyses to identify the active anti-inflammatory compound(s). Ethyl acetate extraction of the supernatant was filtered and fractionated, and a bioactivity-guided isolation approach was employed by testing of resultant fractions for TLR4 and TLR3 suppression (Fig. 2A and B). Cells were activated with LPS or aminoglycoside and then treated with 0.1 mg/ml fraction 2 hours post activation. IFN reporter activity of LPS-treated cells was set at 100% and all other conditions visualized as a percentage of maximum LPS-induced IFN activity (Fig. 2A). Of the 21 fractions, 6 showed high suppressive activity, with IFN reporter activity equivalent to those of untreated cells. Five of these same fractions also reduced TLR3-activation by aminoglycosides (Fig. 2B). We selected the five active fractions that suppressed both TLR4 and TLR3 activation (Fractions #12-16) and 1 control non-active fraction (Fraction #1) and found a clear dose-dependent suppressive effect of active fractions on aminoglycoside-activated cells (Fig. 2C).

Next, the active fractions were combined and purified using a reversed-phase preparative and semi-preparative HPLC, resulting 9 compounds. The isolated compounds were identified via the comprehensive analysis of NMR spectroscopic data and MS data as well as comparison of their NMR spectra with the reported data (Table S1). The majority of compounds isolated all belonged to a single family of β-carboline alkaloids (Fig. 2D). One of the compounds, 1-acetyl-β-carboline (BC5), was previously identified in supernatant of gut lactobacilli strains in a screen for anti-fungal compounds, where it was shown to inhibit fungal filamentation, a virulence associated trait of *Candida albicans* (MacAlpine et al., 2021). However, that study did not test for immunomodulatory activity. We tested each of the compounds for TLR4 inhibitory activity by measuring suppression LPS-induced IFN signaling (Fig. 2E). We found that while almost 65% of the isolated β-carbolines (6/9 compounds) significantly inhibited TLR4 activation, a compound, perlolyrine (BC6) was a standout suppressor, reducing LPS-induced IFN activation by 77 ± 6%. Since these compounds were isolated from the active fractions, we wanted to confirm their activity via independent synthesis. To this end, we selected three compounds, a non-suppressor, flazine (BC1), an intermediate suppressor, 1-furyl-β-carboline-3-carboxylic acid (BC3) and a high suppressor, perlolyrine (BC6) for synthesis.

**Figure 2:**
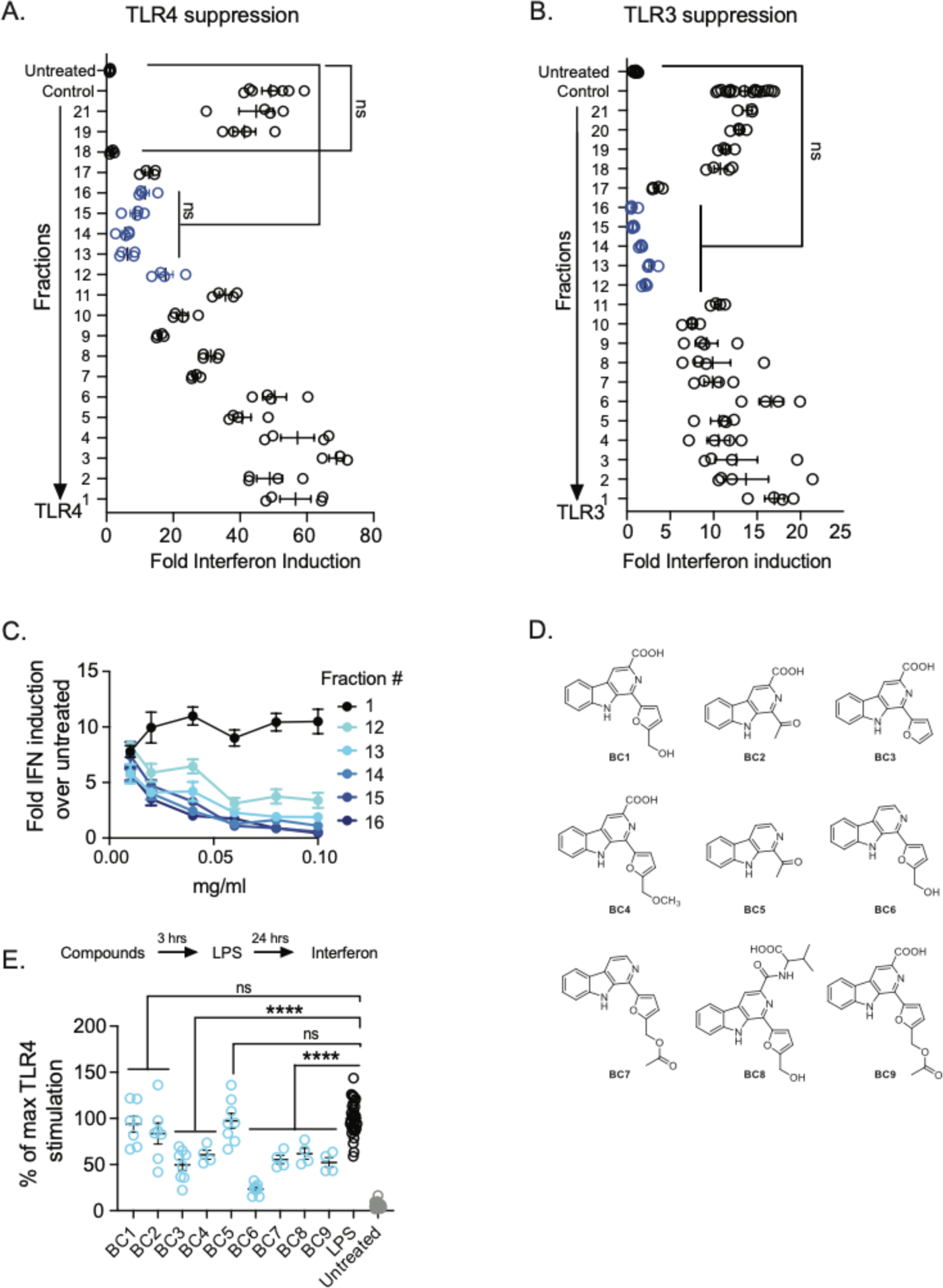
*L. crispatus* produces family of anti-inflammatory β-carboline compounds. Supernatant from *L. crispatus* strain MV-3-USA (HM-636) was fractionated and added to LPS (500 ng/ml) -treated THP1 cells (A) or aminoglycoside (1 mg/ml) - treated THP1 cells (B). Suppressive active fractions common to both treatments are outlined in blue. Active fractions (#12-16) and a control fraction (#1) were added to aminoglycoside-activated cells at indicated doses and interferon reporter activity indicated as fold over untreated cells (C). Structures of β-carboline compounds identified from active fractions (D). THP1 cells were treated with 50 μM each of all 9 β-carboline compounds and then stimulated with LPS for 24 hours (E). Samples were compared using a one-way ANOVA with Dunnett’s correction for multiple comparisons.

We confirmed activity by testing the ability of the independently synthesized compounds to suppress TLR4 activation. As in Figure 3, synthesized BC1 remained non-suppressive. The synthesized BC3 was an intermediate suppressor reducing TLR4 activity to 50.5± 4% of maximum activation when cells were pre-treated with the compound (Fig. 3A). The synthesized BC6 retained the strongest anti-inflammatory activity, reducing LPS-induced interferon signaling to 19.4 ± 1% of maximum TLR4 activity at 100 μM treatment dosage. Of note, this suppressive activity was retained if BC6 was added before or after stimulation after treatment. In contrast, we observed a higher variation in BC3 treatment, cell retained up to 88.4 ± 5.4% of LPS-induced IFN activity when compounds were added after stimulation (Fig. 3B).

Next, we tested if these compounds could also suppress type I IFN signaling downstream of the type I interferon receptor (IFNAR) by treating THP1 cells with IFNβ. We found that cells pretreated with BC6 alone suppressed IFNβ activation to 37± 5% of maximal IFN signaling at higher concentrations of the compound (50,100μM) reducing to 72± 13% of maximal IFN signaling at 10μM doses (Fig. 3C). Similar inhibition levels were observed if the compounds were added after IFNβ stimulation (Fig. 3D). Together we were able to show the β-carboline compounds we identified in bacterial supernatant retained their broad activity profiles after synthesis.

**Figure 3:**
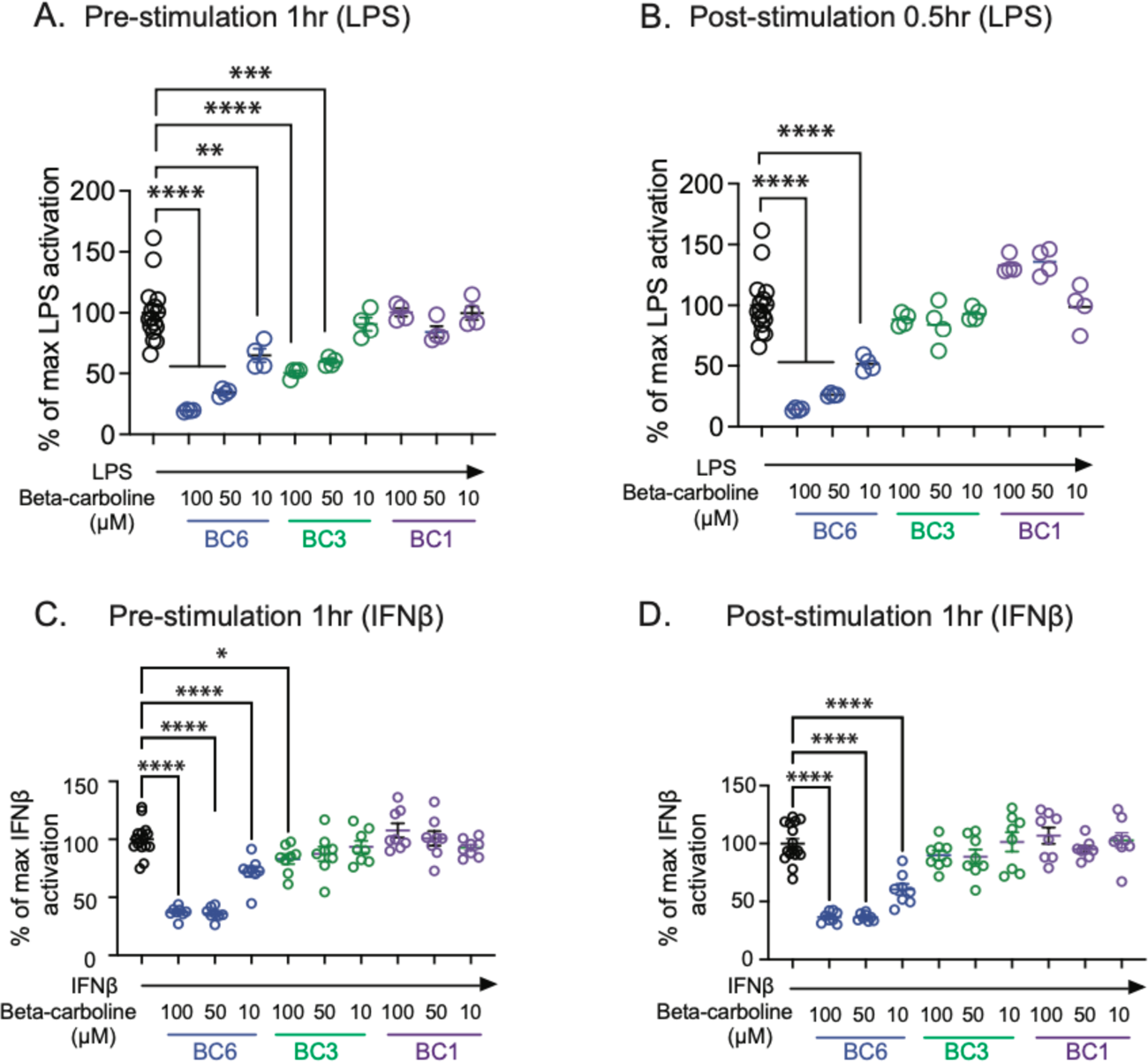
Independently synthesized β-carbolines suppress interferon signaling downstream of TLR4 and IFNAR. THP1 cells were treated with 500 ng/ml LPS (A, B) or 10 ng/ml IFNβ (C, D) and received indicated concentrations of β-carbolines one hour before (A, C) or 0.5-1 hour after stimulation (B, D). ISRE activation was read 24 hours post stimulation. Samples were compared using a one-way ANOVA with Dunnett’s correction for multiple comparisons. Data from 3A, B are representative of 3 independent experiments with 3 biological replicates per experiment. Data from 3B. C are combined from 2 independent experiments with 3-4 biological replicates per condition.

Our experiments to date were conducted on a human macrophage-monocyte reporter cell line and we wanted to verify that these compounds were capable of inhibiting inflammation in primary cells. To do so, we tested the effect of β-carbolines on LPS activation in primary human monocytes. We measured phopho-NFκB staining in CD14+ cells isolated from peripheral blood mononuclear cells (Fig. 4A) and found that BC6 treatment reduced the average frequency of pNFκB+CD14+ cells from 58.8 ± 5.7% to 17.6 ± 4%, slightly lower than the frequency observed in untreated cells, 24± 5.8% (Fig. 4B). Within these cells, the pNFκB mean fluorescence intensity (MFI) was also reduced specifically by BC6 treatment. BC6 but not BC1 or BC3 reduced LPS-mediated 2-fold increase in pNFκB MFI to levels seen in untreated cells (Fig. 4C). Together our data show a robust suppression of NFκB in primary human monocytes.

To determine if this effect was limited to immune cells or could be observed in the epithelia, we employed a mouse vaginal epithelial organoid system (Ali et al., 2020; Ulibarri et al., 2023). We stimulated the organoids with IFNβ and then treated with 100µM BC6. We saw robust induction of multiple interferon-stimulated genes 24 hours post IFNβ stimulation. In contrast to macrophages and monocytes however, BC6 did not suppress IFNAR signaling in epithelial cells (Fig. S3) suggesting that presence of these compounds in the vaginal mucosa may primarily act on innate immune cells.

To determine the transcriptomic effects of these **β-**carbolines, we performed RNA sequencing on human monocytes that received β-carbolines alone or alongside LPS stimulation. We found that BC6 specifically induced transcriptional changes in human monocytes (Fig. 4D). In sharp contrast, BC1 and BC3 did not induce any differentially expressed genes in unstimulated or LPS-stimulated monocytes (Fig. S4A-D). We generated a list of differentially expressed genes induced by BC6 treatment in both the LPS and unstimulated treatment conditions (Table S2). We saw a significant suppression of innate-immunity and inflammation linked genes as identified via gene ontology enrichment (Fig. 4E), with 5 of the 11 categories linked to inflammation and immunity. Importantly, this signature was independent of LPS stimulation suggesting that this anti-inflammatory program is engaged by BC6 prior to stimulation.

**Figure 4:**
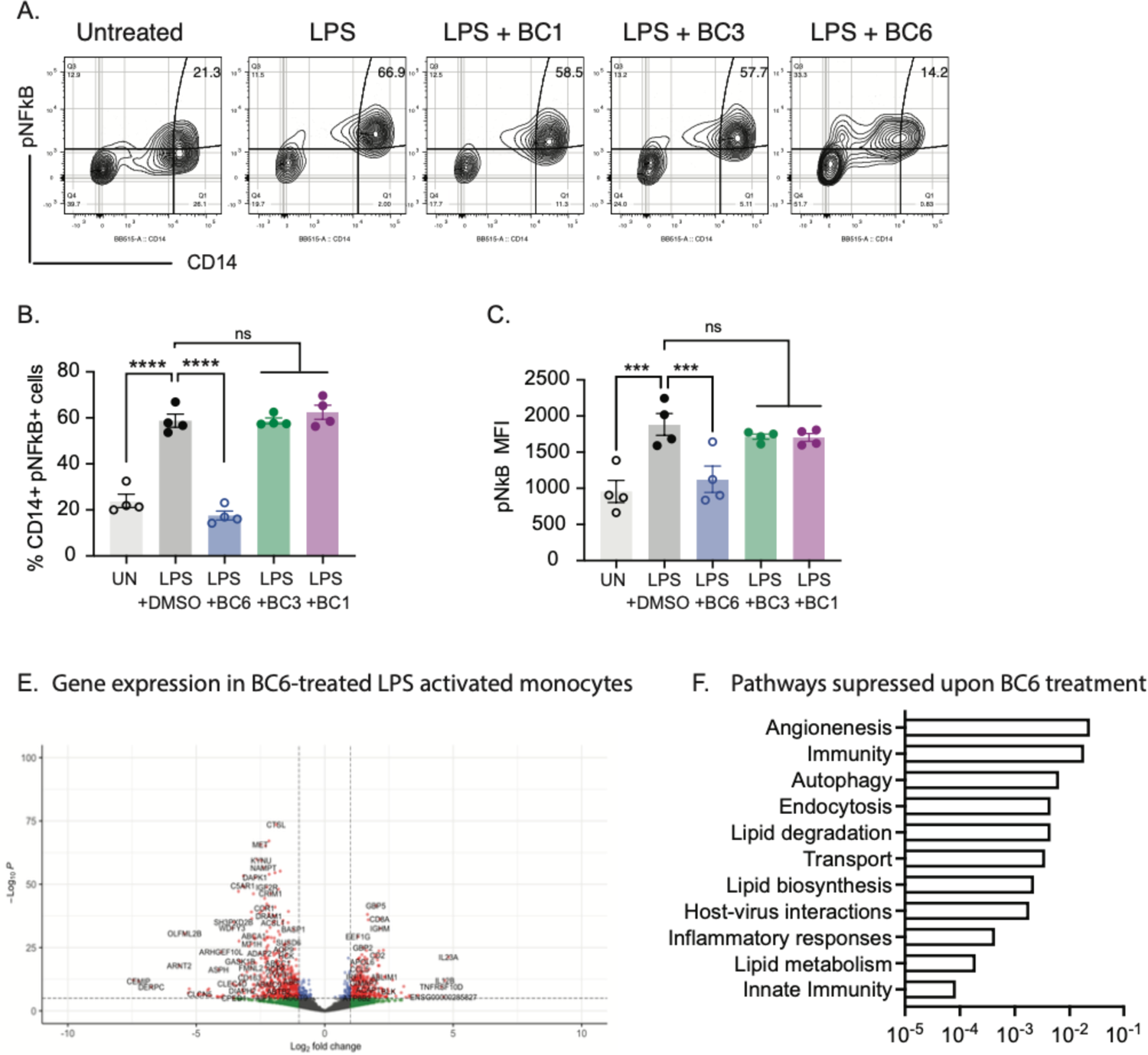
*L. crispatus*-produced β-carbolines suppress inflammatory signaling in primary human monocytes. Primary human monocytes were isolated from peripheral circulating mononuclear cells and treated with 500 ng/ml LPS. One hour post stimulation cells received 10 µM of all three β-carbolines and 24 hours post treatment cells were stained for pNFκB and CD14. Representative FACS plots gated on live cells are shown in (A), pNFκB+CD14+ cells quantified (B) and pNFκB fluorescence intensity quantified in (C). Differentially expressed transcripts upon C6 treatment in LPS-treated monocytes highlighted in volcano plot (E) and significantly enriched pathways in downregulated transcripts shown with adjusted p value (F). Data from 4A-C are representative from 3 independent experiments with 4 replicates per condition.

To test the hypothesis that β-carboline compounds were the mechanism by which lactobacilli suppressed vaginal inflammation, we tested the effect of lactobacilli-supernatant on infection-induced vaginal inflammation using a mouse model of genital herpes infection (Gopinath et al., 2018; Lee et al., 2020). As human vaginal lactobacilli are poor colonizers of the murine vaginal mucosa (Mejia et al., 2023; Vrbanac et al., 2018), we used lactobacilli cell-free supernatant. After intravaginal infection, herpes simplex virus type 2 (HSV-2) undergoes rounds of local mucosal replication in vaginal epithelial cells and vaginal inflammation before traveling to the dorsal root ganglion and causing a range of inflammatory symptoms that can be scored. Unlike in humans, HSV-2 in mice spreads to the central nervous system and results in death of the host, allowing us to track survival. We found that pretreating mice intravaginally with *L. crispatus* supernatant (Fig. 5A) significantly reduced disease scores compared to mice treated with *L. reuteri* supernatant or media controls (Fig. 5B, C). While survival was significantly increased in mice treated with both groups of lactobacilli-supernatant (Fig. 5D), a significant percentage of *L. crispatus* cell-free-supernatant treated mice remained asymptomatic despite equivalent vaginal viral titers (Fig. 5E). These data suggest that vaginal lactobacilli produce effectors that increase disease tolerance.

**Figure 5:**
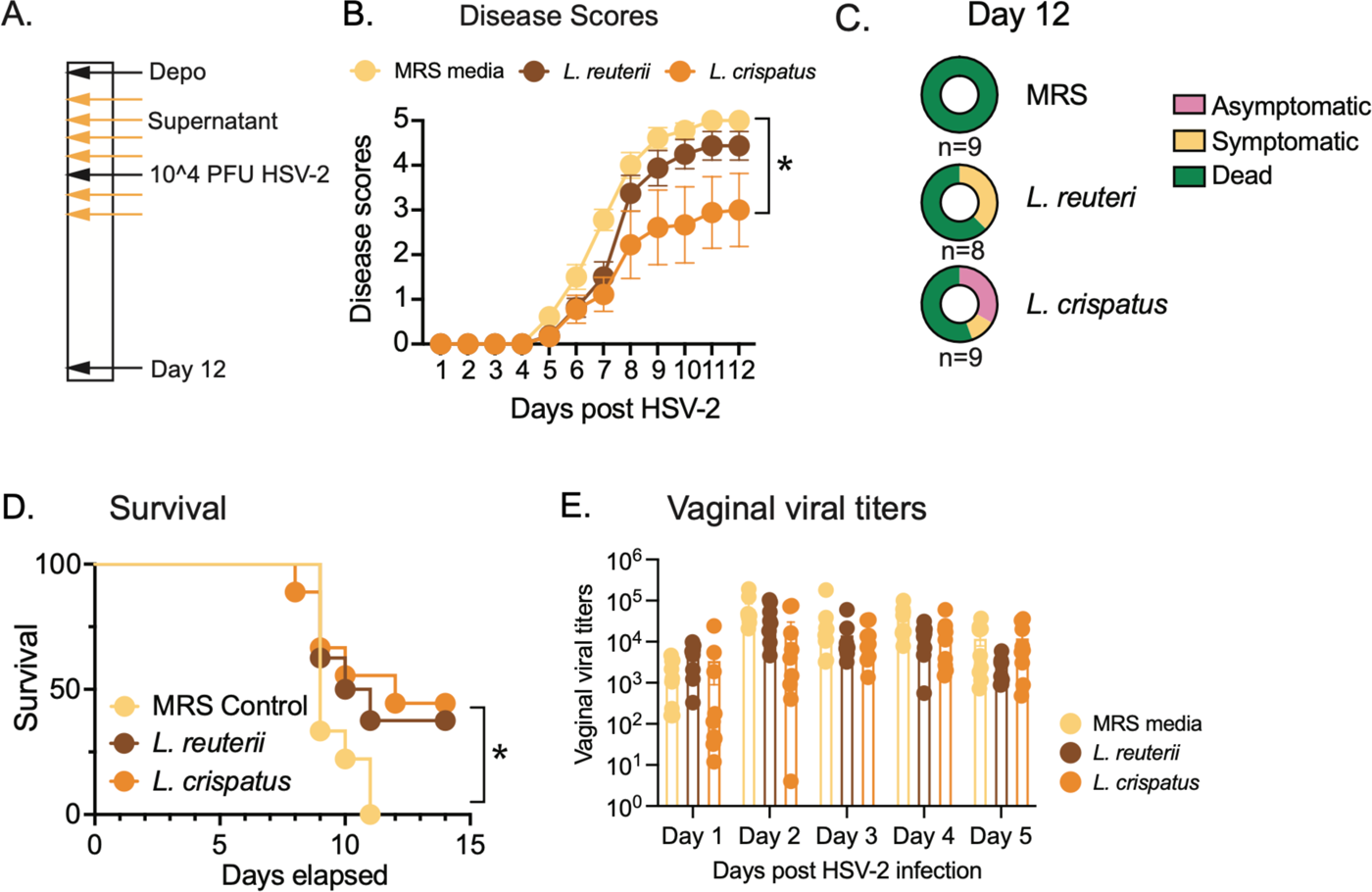
Vaginal lactobacilli supernatant alleviates inflammation during genital herpes infection. Mice were injected with 2 mg medroxyprogesterone and received 10 µl of either cell-free supernatant from *L. crispatus* (HM-637), *L. reuteri* (HM-102) or equivalent volume of media at indicated days (A). Mice were infected with 0.5-1 × 10^4^ PFU HSV-2 and disease scores quantified (B). Day 12 scores were further broken down in (C). Survival was quantified in (D) and infectious virus from vaginal wash quantified in (E). Disease scores were compared using 2-way ANOVA and survival curves were compared using Mantel-Cox test for significance. Data are combined from 2 independent experiments with n = 8-9 mice per condition.

We next wanted to test if BC6 was sufficient to recapitulate the tolerogenic effect of *L. crispatus* supernatant. Since β-carboline compounds had been previously described as being anti-herpetic (Chen et al., 2015), we tested the ability of therapeutic β-carbolines to suppress vaginal inflammation at a time during the disease course when inflammation was known to be driven by the vaginal immune response instead of replicating virus (Lee et al., 2020; Lebratti et al., 2021; Lim et al., 2023). We treated mice intravaginally with 100µM of BC6 daily between days 4-8 post infection (Fig. 6A). Therapeutic intravaginal administration of BC6 was sufficient to significantly reduce disease scores (Fig. 6B, C) while significantly extending survival (Fig. 6D). We confirmed that this was not due to reduced vaginal viral titers as measured post treatment (Fig. 6E). These data confirm that BC6 can recapitulate the tolerogenic anti-inflammatory effect of *L. crispatus* in the context of virus-induced vaginal inflammation.

**Figure 6:**
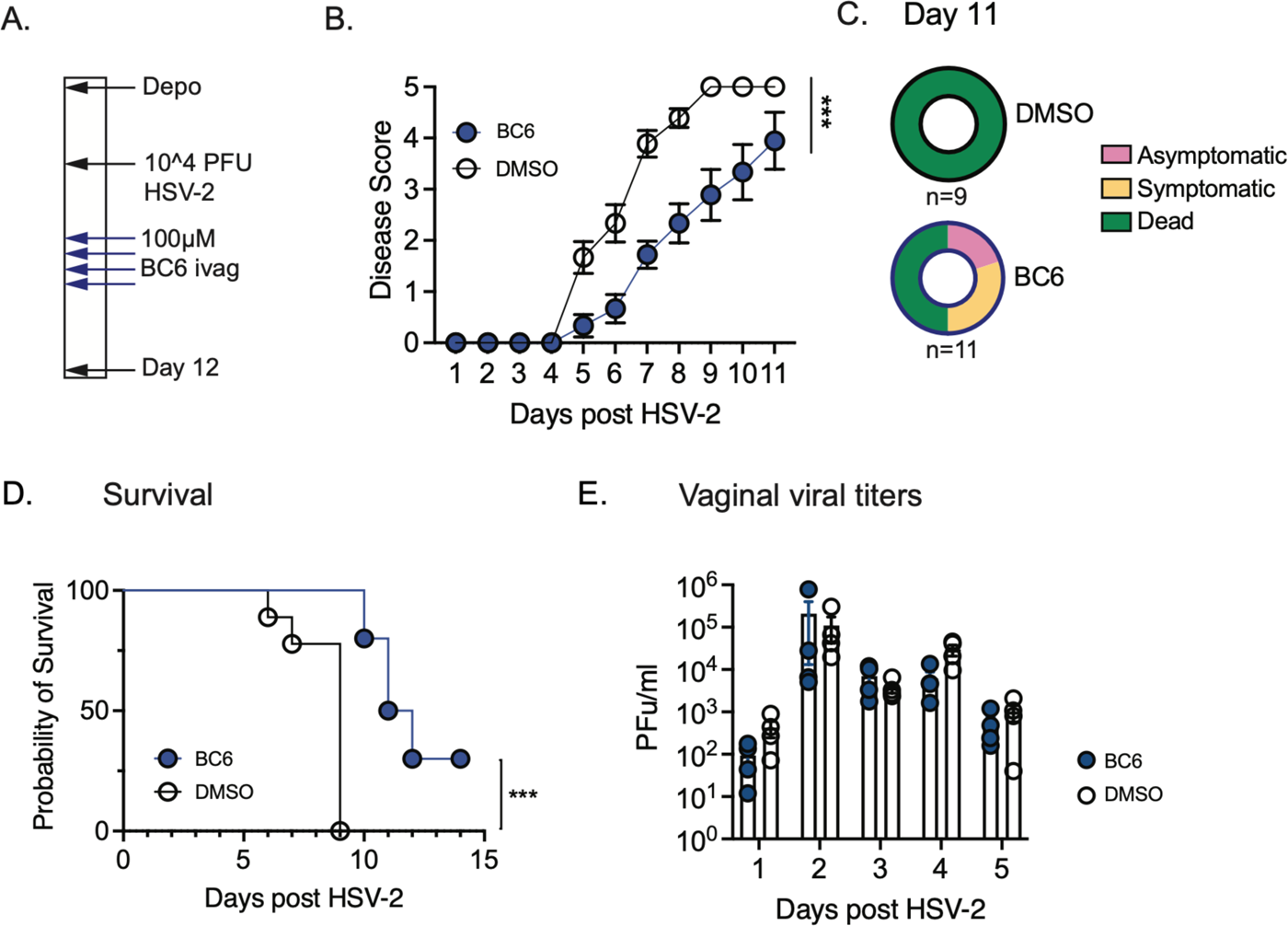
Therapeutic β-carboline application significantly suppresses vaginal inflammation during genital herpes infection. Mice were injected with 2 mg medroxyprogesterone and received 10µl of either β-carboline BC6 or equivalent volume of DMSO (A). Mice were infected with 0.5-1 × 10^4^ PFU HSV-2 and disease scores quantified (B). Day 12 scores were further broken down in (C). Survival was quantified in (D) and infectious virus from vaginal wash quantified in (E). Disease scores were compared using 2-way ANOVA and survival curves were compared using Mantel-Cox test for significance. Data are combined from 2 independent experiments with n = 9-10 mice per condition.

## Discussion

While we have known about the presence of vaginal lactobacilli for almost a century (Thomas, 1928), the mechanisms by which they affect the host, beyond production of lactic acid, remain unknown. We have identified a family of small molecule effectors produced by vaginal lactobacilli, multiple members of which suppress inflammation. The strongest anti-inflammatory compound we identified BC6 has been previously described in the literature as perlolyrine and identified in plant extracts and in food (Herraiz et al., 2023). To our knowledge this is the first example of this compound identified in commensal bacteria. A recent study showed that a gut commensal bacterium *Akkermansia muciniphila* produced a β-carboline harmaline, that mediated anti-inflammatory activity by increasing production of bile acid conjugates (Xie et al., 2023). Other plant-produced β-carbolines have been shown to suppress NFκB signaling (Grodzki et al., 2015; Jin et al., 2022; Niu et al., 2019) however to our knowledge, this is the first report of interferon suppression by β-carbolines.

β-carbolines have been reported to be made by marine and land soil-dwelling bacteria via activity of pictet-splenglerase enzymes (Chen et al., 2018), typically converting L-tryptophan and an aldehyde into β-carboline alkaloids. While ours is the second study to report presence of lactobacilli-produced β-carbolines (MacAlpine et al., 2021), it is the first to identify anti-inflammatory function of a subset of these compounds. The pictet-spenglerase enzymes mediating β-carboline production remain unknown in lactobacilli.

Our results demonstrate broad anti-inflammatory suppression activated by β-carbolines, downstream of multiple pattern recognition receptors and type I interferon receptor suggesting that these compounds may be broadly affecting cell-state. Downregulated genes included multiple members in the innate immune signaling pathway including TLR4-scaffolding protein SASH1(Dauphinee et al., 2013) and large proteins involved in intracellular signaling including multiple members of the dedicator of cytokinesis DOCK1 and DOCK4, which have been shown to exacerbate TLR activation and signaling (Song et al., 2022). The specific downstream signaling pathways activated by BC6 remain unknown. Intriguingly, the anti-inflammatory effect of BC6 seems to be specific to immune cells as BC6 treatment of murine vaginal epithelial organoids did not suppress ISG expression. However, we cannot rule out the possibility that this is due to poor penetration of the β-carboline compounds into the organoids in the matrigel. Given the similarity of the structures of the compounds BC1, BC3 and BC6 (Fig. 2B) we speculate that the differences in suppression may be due as much to accessibility of these compounds to the cytoplasm as compound-specific interactions with host cell proteins. Future work with chemically tagged-β carbolines would help distinguish between these possibilities.

In mice, we found that topical vaginal application of BC6 was sufficient to significantly reduce vaginal inflammation despite equivalent vaginal viral titers in a mouse model of genital HSV-2 infection. In this infection model, both neutrophils and NK cells have been recently found to drive vaginal inflammation 4-8 days post infection (Lim et al., 2023; Lebratti et al., 2021). We found no difference in total neutrophil or NK cell numbers (Fig. S5A, B) in the vaginal lumen; however it is possible that cell signaling within these populations may be altered, leading to a less inflammatory state. Exploring the potential of BC6 as a tolerogenic anti-inflammatory topical compound may be of use in the clinic, especially as it is produced by naturally occurring vaginal lactobacilli strains.

The production of diverse β-carbolines suggest that these compounds may have diverse functions. In support of this, BC5 was previously identified as significantly inhibiting fungal filamentation in *Candida albicans* (MacAlpine et al., 2021). Filamentation is strongly associated with *Candida* pathogenesis suggesting that presence of BC5 β-carboline in the vagina may be a way to maintain *C. albicans* in homeostasis in the vaginal ecosystem. Importantly BC5 showed no anti-inflammatory activity in our study. We speculate that the production of this family of β-carbolines may affect both the vaginal microbial ecosystem and the host to create a more hospitable niche for vaginal lactobacilli. We have identified one strongly anti-inflammatory β-carboline, BC6 or perlolyrine, that can suppress vaginal inflammation when applied topically. Further investigation of vaginal lactobacilli effectors including members of this β-carboline family, would help in the development of novel therapeutics to treat mucosal inflammatory disorders including vaginitis.

## Methods

### Bacterial cultivation and extraction of metabolites

All bacterial strains were obtained from BEI Resources, NIAID, NIH where they were collected as part of the Human Microbiome Project. Strains are identified by strain number whenever possible. Culturing of human vaginal bacteria was performed in an anaerobic chamber (Coy Laboratory Products) with a gas mixture of 5% hydrogen and 10% carbon dioxide unless otherwise stated. Lactobacillus strains with the exception of *L. iners* were grown in De Man–Rogosa–Sharpe (MRS) media. *L. iners* was cultured in NYC III media or in MRS media supplemented with 0.002% L-cysteine and 0.00055% L-glutamine (by volume). All strains were grown from single colonies on MRS agar, cultured for 24 hours in 3 mls and further sub-cultured for 24-48 hours, all under anaerobic conditions at 37 °C. For generation of cell-free media, turbid subcultures were resuspended by vortexing and10 µL from each culture was added to 200 µL of media in triplicate in a 96-well plate (Corning). Plates were sealed with a porous membrane (Diversified Biotech) to facilitate gas exchange and OD600 readings were collected using a plate reader (BioTek) for 48 hours, reading every 30 minutes after a 30 second double orbital shake. At the end of 48 hours, cell cultures were spin filtered over a 22 µM filter and stored at - 80°C for subsequent experiments. Large scale cultivation to isolate metabolites, *L. crispatus* MV-1A-US (HM-637) was inoculated on MRS agar, cultured for 24 hours under anaerobic conditions as described previously. And then, the single colony was transferred into 5 mL of MRS media in 15 mL of tube for 48 hours and further sub-cultured in 1 L of MRS media (total culture volume 14 L). After 3 days, the cultures were centrifuged (8000 r.p.m for 30 minutes at 4 °C) to separate cell pellet from supernatant.

The supernatant was filtered through Whatman qualitative filter paper (grade 3, circle, diameter 125 mm). For metabolite extraction, the filtered supernatant was performed by solvent-partition using ethyl acetate (EtOAc), and the resultant EtOAc solvent mixture containing bacterial metabolites was concentrated on a rotary evaporator. The extraction procedure was repeated three times (total culture volume, 14 L), yielding 5 g of crude extract from the supernatants.

### THP1 culture and experiments

THP1 dual reporter cells (Invivogen, CA) were cultured in complete RPMI (containing 2 mM L-glutamine, supplemented with 10% fetal bovine serum, 1% penicillin-streptomycin, and 25 mM HEPES buffer. All cell culture supplements were purchased from GIBCO. Complete RPMI (cRPMI) was then sterilized by passage through a 0.22 uM vacuum filter. For activation, THP1 cells were plated in a tissue culture-treated 96-well flat bottom plate at a density of 100,000 cells/well. Cells were treated with 50ng/ml, Phorbol 12-myristate 13-acetate (PMA, Invitrogen) for 3 hours. After incubation, cells were centrifuged, PMA-containing media replaced with warm PBS to wash off excess PMA. After a spin wash cycle, THP1 cells in fresh media were placed in the incubator and allowed to rest for 72 hours before experimental use. For bacterial treatment experiments, spent media was discarded, and fresh media with 5% v/v bacterial cell-free supernatant was added to the cells. For stimulation experiments, Pam3Cys2 (20ng/ml, Invivogen), LPS (500ng/ml, Invivogen), Interferon-β (20ng/ml, Peprotech) or Kasugamycin (1mg/ml, Sigma) was added to the cells at indicated time points.

### Bioassay-guided fractionation, purification and identification of β-carboline compounds

The crude extract (5 g) from the supernatants was dissolved in HPLC-grade methanol and filtered through a syringe filter (polytetrafluoroethylene (PTFE), 0.2 µm). The filtered extract was directly injected onto a reversed-phase preparative high-performance liquid chromatography (HPLC) column (Phenomenex Luna C_18_ (2), 250 × 21.2 mm, 5 µm) with a gradient mobile solution (30% methanol/water to 100% methanol for 35 min, 100% methanol isocratic for 10 min; flow rate, 10 ml/min). Fractions were collected every 2 min from 3 min to 45 min, resulting in a total of 21 fractions. The fractions that were able to stimulate anti-inflammatory activity in the human macrophage/monocyte line (THP1s) pre-activated by LPS and aminoglycosides (TLR-4, TLR-3 agonists respectively) were collected in fractions 12-16. The active fractions 12-16 were combined and further fractionated using a reversed-phase preparative HPLC column (Phenomenex Luna C_18_ (2), 250 × 21.2 mm, 5 µm) with a gradient mobile solution (60% methanol/water to 80% methanol/water for 35 min, 80% methanol/water to 100% methanol/water for 1 min, 100% methanol isocratic for 9 min; flow rate, 10 ml/min). Sub-fractions were collected every 2 min from 5 min to 45 min, resulting in a total of 20 fractions. According to the LC/MS analysis for the sub-fractions, sub-fraction 5 (15 mg) was then subjected to reversed-phase semi-preparative HPLC (Phenomenex Luna C_18_ (2), 250 × 10 mm, 5 µm) using the following gradient solvent system: 38% acetonitrile/water isocratic for 40 min; gradient to 45% acetonitrile/water for 10 min; then 45% acetonitrile/water to 50% acetonitrile/water for 10 min, 50% acetonitrile/water to 100% acetonitrile/water for 1 min, and 100% acetonitrile/water isocratic for 9 min; flow rate, 2 ml/min) to isolate BC1 (*t_R_* = 20 min, 1.2 mg), BC2 (*t_R_* = 32 min, 3.0 mg), BC3 (*t_R_* = 44 min, 0.7 mg), and BC4 (*t_R_* = 48 min, 0.9 mg). Sub-fraction 7 (25 mg) was subjected to reversed-phase semi-preparative HPLC (Phenomenex Luna C_18_ (2), 250 × 10 mm, 5 µm) using the following gradient solvent system: gradient 30% acetonitrile/water to 50% acetonitrile/water for 40 min; 50% acetonitrile/water to 100% acetonitrile/water for 10 min; 100% acetonitrile/water isocratic for 10 min; flow rate, 2 ml/min) to isolate BC5 (*t_R_* = 46 min, 1.1 mg). Sub-fraction 8 (67 mg) was purified by reversed-phase semi-preparative HPLC (Phenomenex Luna C_18_ (2), 250 × 10 mm, 5 µm) using the following gradient solvent system: gradient 30% acetonitrile/water to 70% acetonitrile/water for 40 min; 75% acetonitrile/water to 100% acetonitrile/water for 10 min; 100% acetonitrile/water isocratic for 10 min; flow rate, 2 ml/min) to isolate BC6 (*t_R_* = 38 min, 2.0 mg), BC7 (*t_R_* = 41 min, 3.2 mg), BC8 (*t_R_* = 43 min, 0.9 mg), and BC9 (*t_R_* = 45 min, 1.4mg).

The structures of compounds found in the active fractions were determined by the comprehensive analysis of ^1^H NMR spectroscopic data (Tables S1 and S2) and high-resolution MS data as well as comparison of their NMR spectra with the reported data(Li et al., 2011; Byun et al., 2023; Tang et al., 2008; Cheng, Y et al.; Shin, Hae Jae et al., 2010; Fuda et al., 2019) The structure of BC8 was elucidated via the comprehensive analysis of ^1^H, ^13^C and two-dimensional (2D) NMR spectroscopic data and high-resolution MS data. Accordingly, the isolated compounds were identified as flazin (BC1), 1-acetyl-3-carboxy-β-carboline (BC2), 1-furyl-β-carboline-3-carboxylic acid (BC3), 1-[5-(methoxymethyl)-2-furanyl]-β-carboline-3-carboxylic acid (BC4), 1-acetyl-β-carboline (BC5), perloyrine (BC6), and *O*-Acetylperlolyrine (BC7), and among the isolated compounds, compounds BC8 and BC9 were unreported compounds, which named flazinylvaline (BC8), and acetylflazin (BC9), respectively.

Flazin (BC1): amorphous yellow solid; ^1^H NMR (600 MHz, CD_3_OD) see Table S1; ESIMS *m*/*z* 309.1 [M + H]^+^.

1-Acetyl-3-carboxy-β-carboline (BC2): amorphous yellow solid; ^1^H NMR (600 MHz, CD_3_OD) see Table S1; ESIMS *m*/*z* 255.1 [M + H]^+^.

1-Furyl-β-carboline-3-carboxylic acid (BC3): amorphous yellow solid; ^1^H NMR (600 MHz, CD_3_OD) see Table S1; ESIMS *m*/*z* 279.1 [M + H]^+^.

1-[5-(Methoxymethyl)-2-furanyl]-β-carboline-3-carboxylic acid (BC4): amorphous yellow solid; ^1^H NMR (600 MHz, CD_3_OD) see Table S1; ESIMS *m*/*z* 323.1[M + H]^+^.

1-Acetyl-β-carboline (BC5): amorphous yellow solid; ^1^H NMR (600 MHz, CD_3_OD) see Table S1; ESIMS *m*/*z* 211.2 [M + H]^+^.

Perloyrine (BC6): amorphous yellow solid; ^1^H NMR (600 MHz, CD_3_OD) see Table S1; ESIMS *m*/*z* 265.2 [M + H]^+^.

*O*-Acetylperlolyrine (BC7): amorphous brown solid; ^1^H NMR (600 MHz, CD_3_OD) see Table S1; ESIMS *m*/*z* 307.2 [M + H]^+^.

Flazinylvaline (BC8): amorphous brown solid; ^1^H NMR (600 MHz, CD_3_OD) and ^13^C NMR (125 MHz, CD_3_OD) see Table S2; ESIMS *m*/*z* 408.2 [M + H]^+^; HRESIMS *m*/*z* 408.1553 [M + H]^+^ (calcd. for C_22_H_22_N_3_O_5_, 408.1554).

Acetylflazin (BC9): amorphous brown solid; ^1^H NMR (600 MHz, CD_3_OD) and ^13^C NMR (125 MHz, CD_3_OD) see Table S2; ESIMS *m*/*z* 351.1 [M + H]^+^; HRESIMS *m*/*z* 351.0977 [M + H]^+^ (calcd. for C_19_H_15_N_2_O_5_, 351.0975).

### NMR spectroscopy

Nuclear magnetic resonance (NMR) spectroscopic data were obtained using Bruker NEO NMR system (^1^H: 600 MHz, ^13^C: 150 MHz) (Bruker, Billerica, MA) with CD_3_OD (Cambridge Isotope Laboratories, Inc., Tewksbury, MA). Chemical shifts are presented on a *δ* scale. Residual protium in the NMR solvent (*δ*_H_: 3.31, *δ*_C_: 49.15) was used as the reference of chemical shifts. Data are represented as follows: assignment, chemical shift, integration, multiplicity (s, singlet; d, doublet; t, triplet; m, multiplet) and coupling constant in Hz. Full assignment of protons and carbons were completed on the basis of the following 2D NMR spectroscopy experiments: gradient ^1^H–^1^H correlation spectroscopy, gradient ^1^H–^13^C heteronuclear single quantum coherence, gradient ^1^H–^13^C heteronuclear multiple bond connectivity. The software Mnova was used to analyze NMR data.

### High-resolution mass spectrometry for β-carboline compounds

High-resolution electrospray ionization mass spectrometry (HRESIMS) data were collected using Agilent MassHunter Work Station LC/MS Data Acquisition 10.1 and Agilent LC-QTOF Mass Spectrometer 6530 (Agilent Technologies, Santa Clara, CA) equipped with an Agilent 1290 uHPLC system and electrospray ionization detector scanning from *m/z* 50 to 3,200. Samples were injected into a reverse-phase column (Kinetex EVO C18, 5 μm, 100 × 4.6 mm, flow rate 0.4 mL/min) with a gradient of H_2_O (A)-Acetonitrile (B): 0–15 min, 10–100% B; 15–16 min, 100-10% B; 16–21min, 10% B. Agilent MassHunter Qualitative Analysis B.07.00 software was used to analyze the data.

### Mouse genital herpes infections

6-8 week old C57BL/6 female mice were obtained from Charles River Laboratories (strain 027) and injected subcutaneously with 2mgs medroxyprogesterone (Depo Provera, List Biologicals). Five days after injection, mice were infected intravaginally with 5000 PFU HSV-2 (186syn+ strain, a kind gift from Dr. David Knipe). Vaginal lavage was collected by swabbing mice with a calginate swab (Puritan) and pipetting 50µL sterile PBS into the vaginal canal, 5-10 times. Mice were scored for disease symptoms using the following metric: vaginal erythema and inflammation without hair loss as scored as 1, hair loss in the perianal area scored as 2, morbility symptoms including hunched posture, ruffled fur due to lack of grooming scored as 3, hind limb paralysis scored as 4 while death either by disease or via euthanasia due to weight loss in excess of 20% of initial weight, or inability to access food due to hind limb paralysis on both legs was scored as 5. Vaginal viral titers were quantified via plaque assay on Vero cells and virus was passaged on Vero cells as well.

### Human monocyte experiments

Human monocytes were isolated using CD14 positive magnetic selection (Stemcell, Canada), rested for 24 hours and treated with LPS and indicated β-carbolines. β-carbolines were synthesized by WuXi AppTec (Shanghai, China). Cells were collected for flow cytometry and RNA sequencing experiments 24 hours post stimulation and treatment. For flow cytometry experiments, cells were stained 24 hours post stimulation with fixable live/dead, anti-pNFκB and CD14 antibodies and run on a BD Fortessa flow cytometer. For RNA sequencing experiments, all samples were treated in triplicate and libraries were generated using the Kapa mRNA HyperPrep Kit, and they were sequenced on a NextSeq1000/2000 using NextSeq 1000/2000 P2 Reagents (200 Cycles) with dual indexing. . Adapters and low-quality bases were trimmed from paired-end raw reads using Trimmomatic (v0.39), followed by alignment to the human genome (GRCh38) using STAR aligner (v2.7.10b). Gene expression was quantified using RSEM (v1.2.29) to generate read counts. Read counts were analyzed using DESeq2(Love et al., 2014) and gene expression pathways analyzed using DAVID(Huang et al., 2009) and cluster profiler (Yu et al., 2012).

### Murine vaginal epithelial organoid experiments

Vaginal organoids frozen at passage #3 were a kind gift from Dr. Lalit Beura and isolated as previously described (Ulibarri et al., 2023). Organoids were thawed, plated in Matrigel (Corning) and passaged every 7-10 days. Organoid culture media was supplemented every 2 days with minor changes. Complete media was DMEM/F12 supplemented with B27 (20 ul/mL), Amphotericin B (0.25ug/mL), TGFβ-inhibitor (0.5uM) and EGF (100 ng/mL).

ROCK inhibitor (10uM) was included for the first four days of culture after each passage. All supplements were obtained from Peprotech (New Jersey, USA). except ROCK inhibitor which was purchased from Tocris (Minneapolis, USA)For passaging, matrigel domes containing the organoids were liquified and resuspended in ice-cold PBS. Organoid cells were dissociated using 0.25% Trypsin in a 37°C water bath for 5 minutes. After incubation, DMEM/F12 medium with 10% fetal bovine serum (FBS) was added to halt trypsinization. The cell pellet was resuspended in complete DMEM/F12 media to remove residual FBS. Finally, cells were resuspended in Matrigel by gentle pipetting and reseeded 1:4 in a 24 well plate using 20uL Matrigel domes. Matrigel domes were solidified for 1-hour, complete DMEM/F12 media was added to the organoids. For gene expression experiments, RNA was isolated from organoid pellets and gene expression quantified via qPCR using previously published primers (Gopinath et al., 2018).

## Acknowledgments

We thank Dr. Wendy Garrett and members of the Garrett lab for useful discussion and feedback. We thank Dr. Curtis Smith, Dr. Anu Natarajan and Dr. Jinbo Lee for consultation on β-carboline synthesis and oversight of contract research organization orders at WuXi AppTec. We thank Dr. David Knipe for generous gift of HSV-2 viral strains and fruitful discussions.

## Funding

This project was funded by Searle Fellowship and Blavatnik Biomedical Accelerator awarded to SG and National Science Fellowship awarded to VJG. This work was supported by a National Research Foundation of Korea (NRF) grant funded by the Korean government (MSIT; grant numbers 2019R1A5A2027340 and 2021R1A2C2007937) awarded to KHK.

## Supplementary Figures

**Table S1.** NMR data of compounds BC1-BC7 (*δ* ppm).*^a^*.

| Position | BC1<br>(Li et al.,<br>2011) | BC2<br>(Byun et al.,<br>2023) | BC3<br>(Tang et al.,<br>2008) | BC4<br>(Cheng, Y et<br>al.) | BC5<br>(Shin, Hae Jae<br>et al., 2010) | BC6<br>(Li et al.,<br>2011) | BC7<br>(Fuda et al.,<br>2019) |
| --- | --- | --- | --- | --- | --- | --- | --- |
| | $\delta_H$ (J in Hz) | $\delta_H$ (J in Hz) | $\delta_H$ (J in Hz) | $\delta_H$ (J in Hz) | $\delta_H$ (J in Hz) | $\delta_H$ (J in Hz) | $\delta_H$ (J in Hz) |
| 3 | - | - | - | - | 8.31, d (5.4) | 8.02, d (5.4) | 8.07, d (4.8) |
| 4 | 8.55, s | 8.55, s | 8.55, s | 8.55, s | 8.46, d (4.8) | 8.29, d (4.8) | 8.32, d (4.8) |
| 5 | 8.23, d (7.8) | 8.25, d (7.8) | 8.22, d (7.8) | 8.23, d (7.8) | 8.22, d (7.8) | 8.19, d (7.8) | 8.21, d (8.4) |
| 6 | 7.30, t (7.2) | 7.33, t (7.2) | 7.30, t (7.2) | 7.31, td (7.2,<br>1.2) | 7.31, t (7.8) | 7.29, t (7.8) | 7.31, td (7.2,<br>0.6) |
| 7 | 7.58, t (7.2) | 7.59, td (7.2,<br>1.2) | 7.57, m | 7.58, td (7.2,<br>1.2) | 7.59, td (8.4,<br>1.2) | 7.59, td (7.8,<br>1.2) | 7.60, m |
| 8 | 7.72, d (7.8) | 7.71, t (7.2) | 7.72, d (7.8) | 7.73, d (7.8) | 7.71, d (8.4) | 7.71, d (8.4) | 7.75, d (7.8) |
| 2' | - | 2.94, s | - | - | 2.83, s | - | - |
| 3' | 7.53, d (3.0) | - | 7.57, m | 7.54, d (3.0) | - | 7.23, d (3.6) | 7.27, d (3.6) |
| 4' | 6.58, d (3.0) | - | 6.71, dd (3.6,<br>1.8) | 6.69, d (3.6) | - | 6.60, d (3.0) | 6.75, d (3.6) |
| 5' | - | - | 7.58, br s | - | - | - | - |
| 6' | 4.77, s | - | - | 4.69, s | - | 4.77, s | 5.35, s |
| 6'-OCH<sub>3</sub> | - | - | - | 3.47, s | - | - | - |
| 2'' | - | - | - | - | - | - | 2.14, s |

<sup>a</sup> Measured at 600 MHz in CD<sub>3</sub>OD; Coupling constants (Hz) are given in parentheses.

**Table S2: Differentially regulated transcripts in BC-6 treated monocytes.** Table is available as a supplementary excel sheet.

**Supplementary Figure 1:**
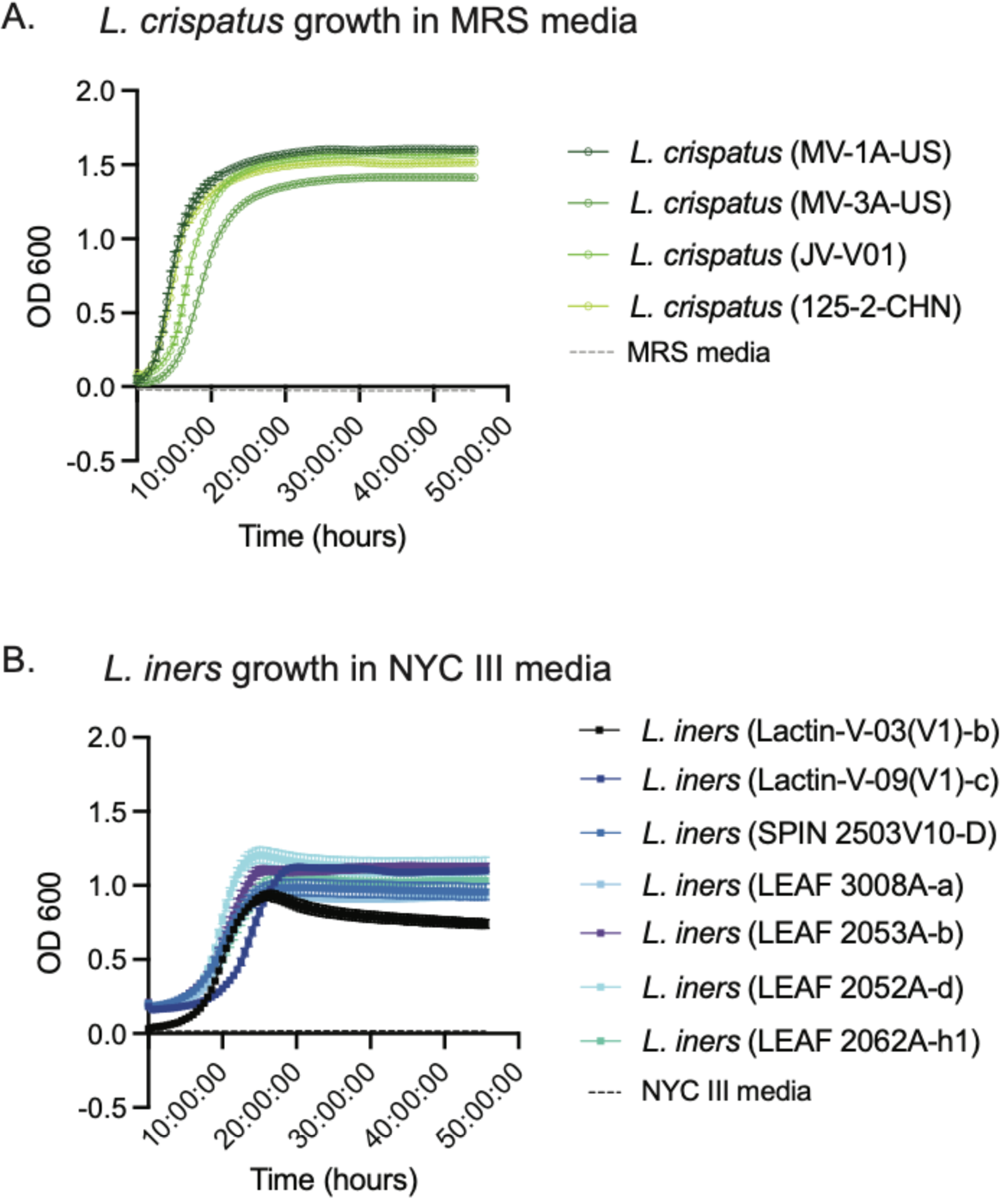
Lactobacilli strains growth curve show maximized growth by 48 hours. Indicated *L. crispatus* strains (A) and *L. iners* strains (B) were sub-cultured in MRS and NYC media respectively and growth monitored via absorbance reading at OD600 for 48 hours. All strains show maximal growth by 48 hours.

**Supplementary Figure 2:**
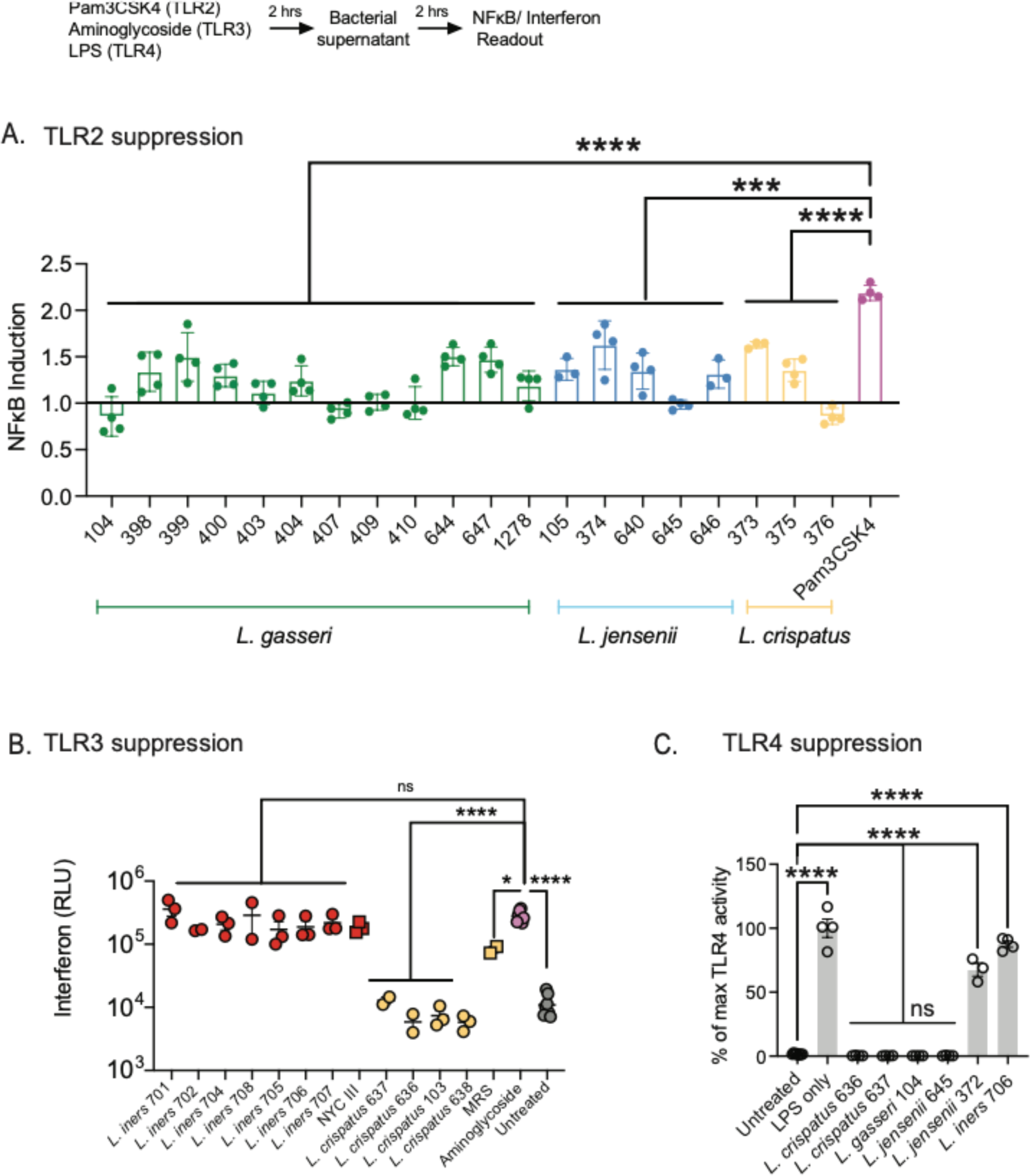
Suppression of cell surface and endosomal activated inflammatory responses observed across multiple strains from *L. crispatus*, *L. gasseri* and *L. jensenii* species. Indicated bacterial strains numbered with the Human Microbiome Project strain numbers, were grown from single colonies for 28 hours in MRS media and filtered-cell-free supernatant added to THP1 reporter monocytes at 5% v/v and NFκB (A) or interferon (B) was read 24 hours after addition. THP1 cells were stimulated with 20 ng/ml PAM3Cys to activate TLR2 responses (A), 1 mg/ml kasugamycin to activate TLR3 responses (B) or 500 ng/ml LPS to activate TLR4 responses (C). Samples were compared using a one-way ANOVA with Dunnett’s correction for multiple comparisons.

**Supplementary Figure 3:**
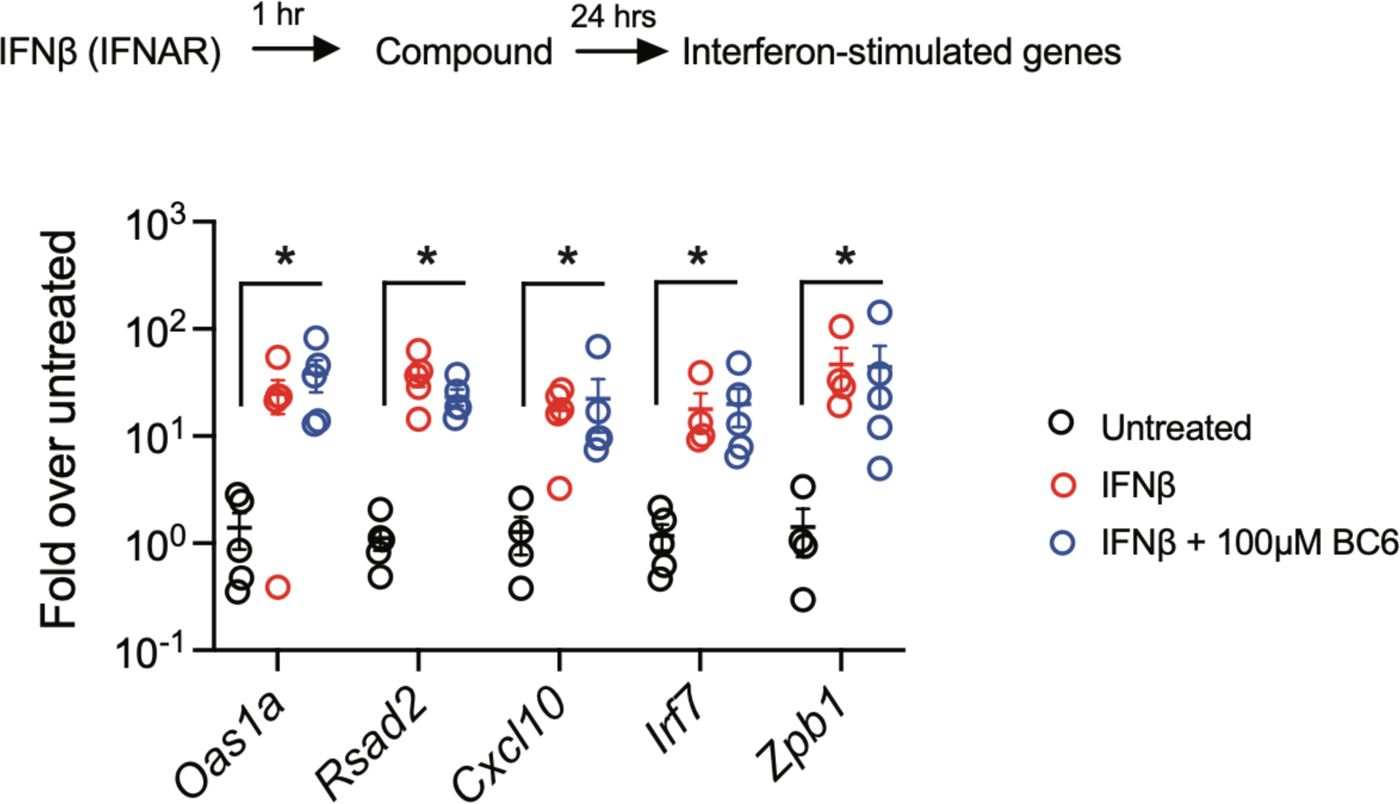
BC6 does not suppress interferon responses. Vaginal epithelial organoids were stimulated with 10 ng/ml IFNβ and then treated with 100 µM BC6. RNA was collected 24 hours post stimulation and gene expression of indicated interferon-stimulated genes quantified via qPCR. Conditions were compared via Mann-Whitney test, using Holm-Sidak correction for multiple comparisons.

**Supplementary Figure 4:**
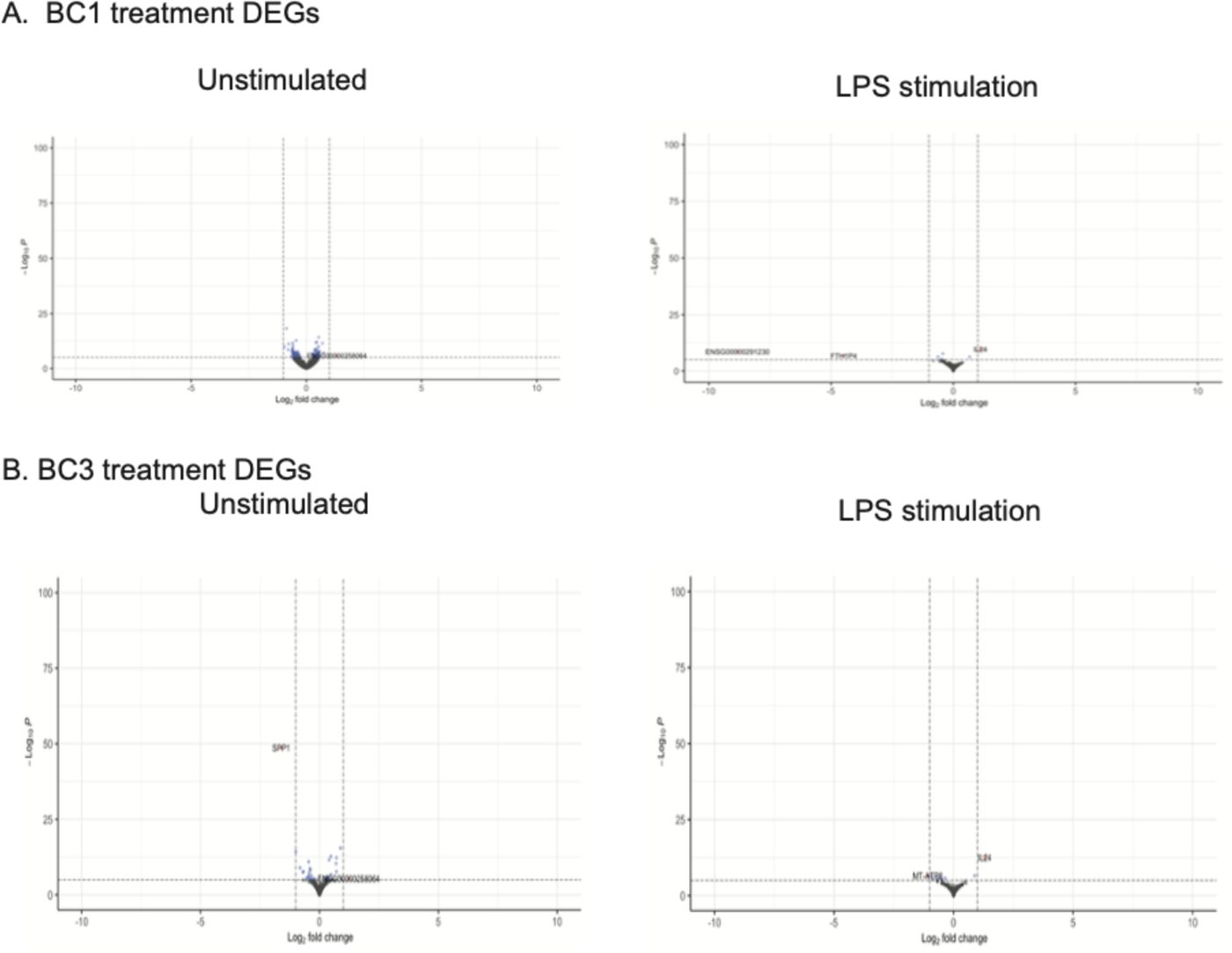
Differentially expressed transcripts upon BC1 and BC3 treatment. Primary human monocytes were isolated from peripheral circulating mononuclear cells and treated with 500 ng/ml LPS or left unstimulated. Volcano plots show differentially expressed transcripts upon BC1 (A) and BC3 (B) treatment in unstimulated and LPS-stimulated cells.

**Supplementary Figure 5:**
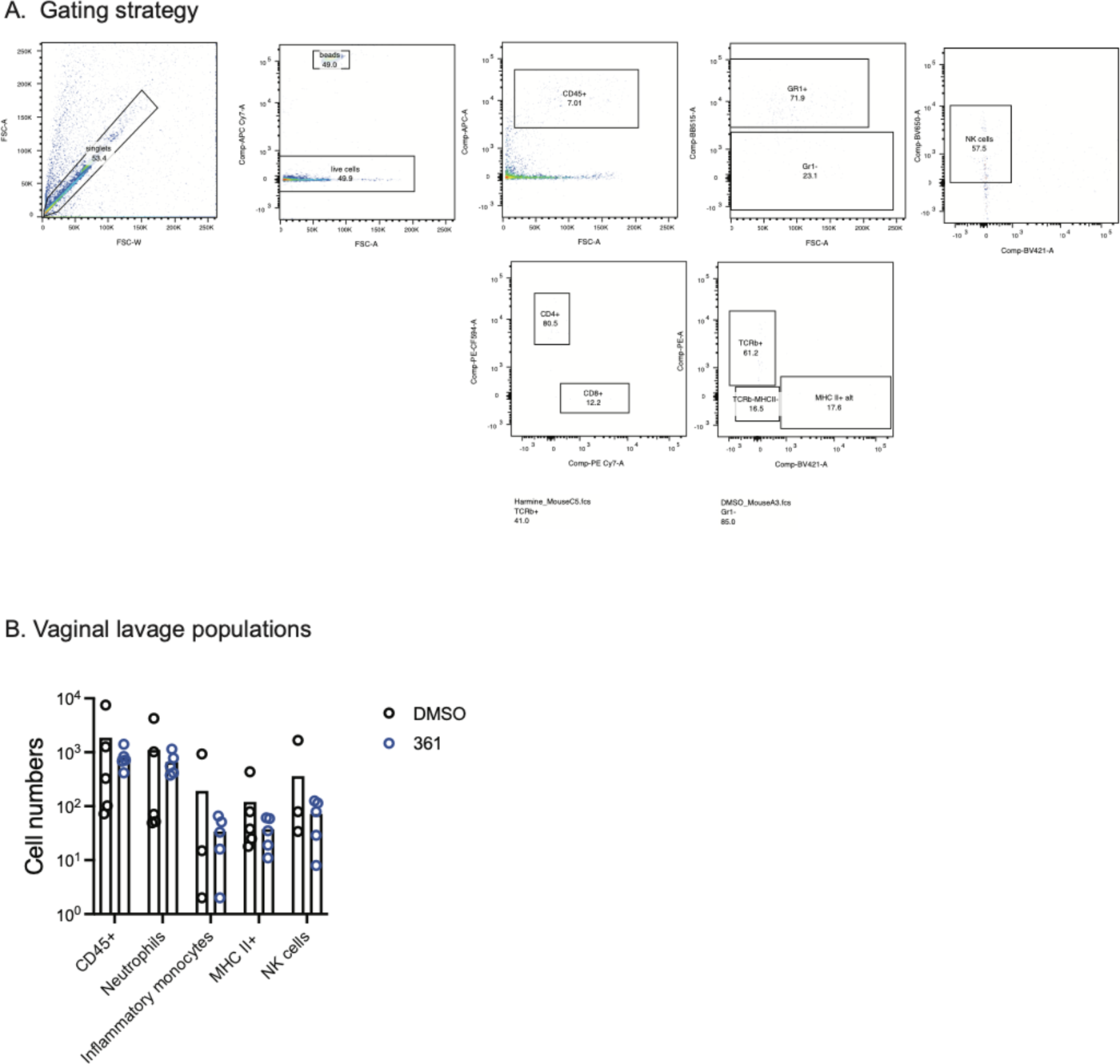
Vaginal luminal cell populations after BC6 treatment in HSV-2 infected mice. Vaginal lavage was collected on day 7 post HSV-2 infection from DMSO and BC6 treated mice. Cell pellets were stained, and populations gated as shown in (A) and quantified in (B).

## Notes

### Competing Interest Statement

C.W., M.M, S.B, J.C., K.H.K. and S.G. are co-inventors on a patent related to this work.

